# Unequivocal mapping of molecular ether lipid species by LC-MS/MS in plasmalogen-deficient mice

**DOI:** 10.1101/2020.04.29.066530

**Authors:** Jakob Koch, Katharina Lackner, Yvonne Wohlfarter, Sabrina Sailer, Johannes Zschocke, Ernst R. Werner, Katrin Watschinger, Markus A. Keller

**Affiliations:** Institute of Human Genetics, Medical University of Innsbruck, 6020 Innsbruck, Austria; Institute of Biological Chemistry, Biocenter, Medical University of Innsbruck, 6020 Innsbruck, Austria

## Abstract

Deficient ether lipid biosynthesis in rhizomelic chondrodysplasia punctata and other disorders is associated with a wide range of severe symptoms including small stature with proximal shortening of the limbs, contractures, facial dysmorphism, congenital cataracts, ichthyosis, spasticity, microcephaly, and mental disability. Mouse models are available but show less severe symptoms. In both humans and mice it has remained elusive which of the symptoms can be attributed to lack of plasmanyl or plasmenyl ether lipids. The latter compounds, better known as plasmalogens, harbor a vinyl ether double bond conferring special chemical and physical properties. Discrimination between plasmanyl and plasmenyl ether lipids is a major analytical challenge, especially in complex lipid extracts with many isobaric species. Consequently, these lipids are often neglected also in recent lipidomic studies. Here we present a comprehensive LC-MS/MS based approach that allows unequivocal distinction of these two lipid subclasses based on their chromatographic properties. The method was validated using a novel plasmalogen-deficient mouse model which lacks plasmanylethanolamine desaturase and therefore cannot form plasmenyl ether lipids. We demonstrate that plasmanylethanolamine desaturase deficiency causes an accumulation of plasmanyl species, a little studied but biologically important substance class.

## Introduction

In mammalia, more than 20% of all glycerophospholipids are considered to be ether lipids ^1^. These lipids carry an ether-linked fatty alcohol rather than an ester-linked fatty acid at their *sn-*1 position and occur abundantly within the lipid classes of phosphatidylethanolamines (PE) and phosphatidylcholines (PC). Plasmalogens are a subgroup of ether lipids and defined by a characteristic 1-O-alk-1’-enyl ether (vinyl ether) double bond (Figure 1A). Although their exact functional range still has to be fully uncovered, they have been shown to be involved in shaping membrane properties, to act as potent antioxidants, and to be involved in inflammatory signal transduction ^2^.

**Figure 1:**
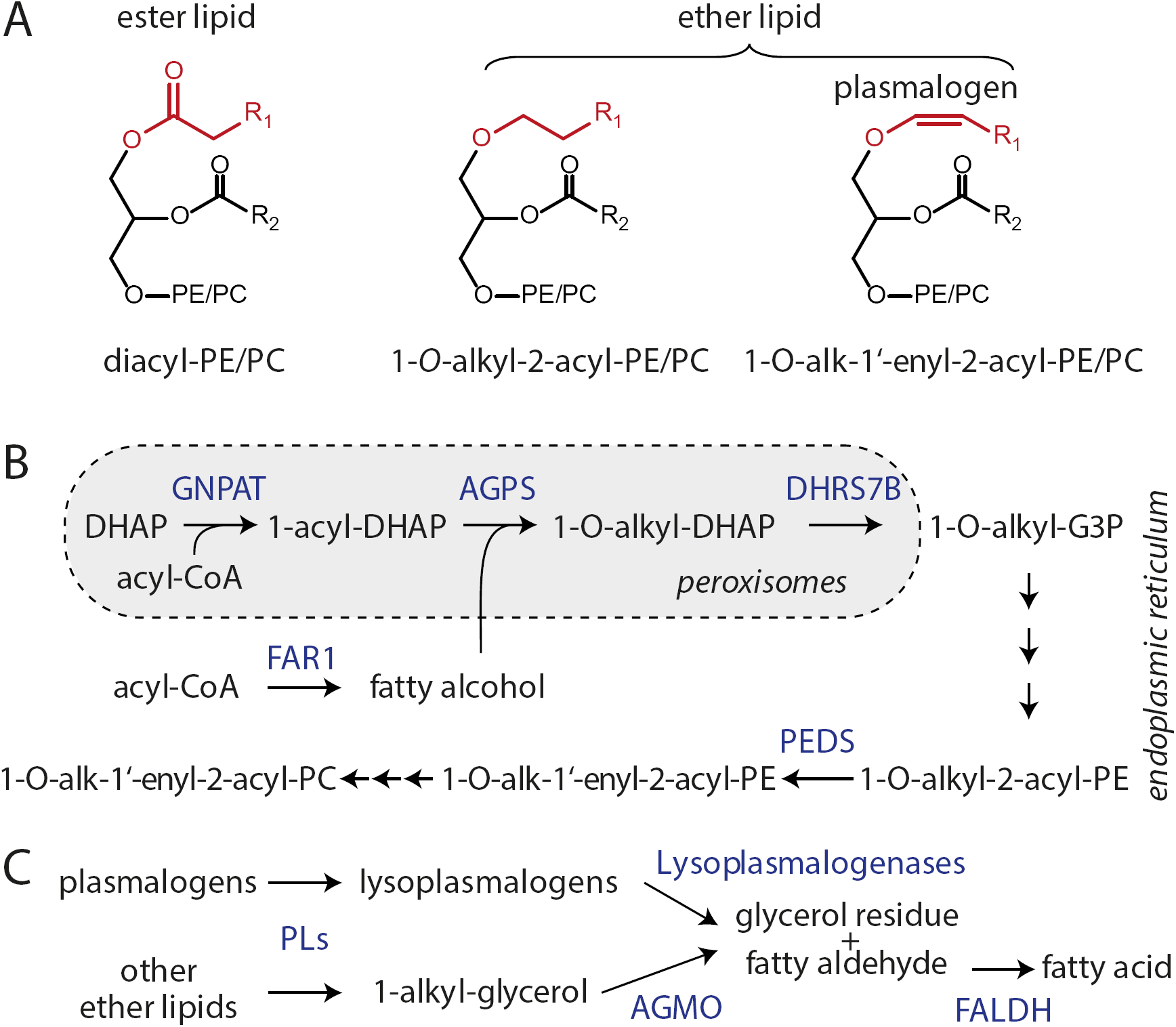
Ether lipid and plasmalogen metabolism. A) Structures of diacyl-phosphatidylethanolamine (PE) and phosphatidylcholine (PC) ester lipids, as well as ether lipids characterized by 1-O-alkyl and 1-O-alk-1’-enyl (=plasmalogens) substitution at the sn-1 position. B) Biosynthesis of ether lipids and plasmalogens crucially relies on functional peroxisomes and is dependent on fatty alcohol availability, which is controlled by a fatty acid reductase (FAR1). Plasmalogens are then formed in the endoplasmic reticulum by plasmanylethanolamine desaturase (PEDS). Further enzymes are: GNPAT: glycerone phosphate O-acyltransferase; AGPS: alkylglycerone phosphate synthase; DHRS7B: alkyl/acyl DHAP reductase. Substrates: G3P: glyceraldehyde 3-phosphate; DHAP: dihydroxyacetone phosphate; CoA: coenzyme A. C) Degradation routes of ether lipids require prior cleavage of sn-2 residues and/or polar headgroup by phospholipases (PLs). 1-O-alkyl-glycerols are cleaved by alkylglycerol monooxygenase (AGMO), while plasmalogenases degrade plasmalogens. Both processes produce fatty aldehydes that are cleared by fatty aldehyde dehydrogenase (FALDH).

The biosynthesis of ether lipid starts with the synthesis of 1-O-alkyl-G3P in peroxisomes (Figure 1B) and is impaired in specific inherited enzyme/transporter deficiencies (rhizomelic chondrodysplasia punctate) as well as peroxisome biogenesis disorders (Zellweger spectrum disorders) ^3^. Plasmalogens are subsequently formed in the endoplasmic reticulum from 1-O-alkyl-PE precursors by the action of the enzyme plasmanylethanolamine desaturase (PEDS) which was only recently identified to be encoded by *TMEM189* ^4,5^. Some neurodegenerative conditions, including Alzheimer’s disease, are thought to be associated with impaired plasmalogen homeostasis ^6^. In contrast to ester-linked acyl side chains, alkyl and alk-1’-enyl residues cannot be readily remodeled, instead these lipids have to be fully degraded by dedicated enzymatic routes (Figure 1C) ^7–9^.

A well-known problem for lipidomic analysis is the differentiation of isobaric phospholipid species, especially among ether lipids ^10,11^. While alkyl and acyl species can be differentiated using a high mass resolution, isobaric 1-O-alkyl (plasmanyl) and 1-O-alk-1’-enyl (plasmenyl) lipids cannot be distinguished by their exact masses and fragmentation patterns alone. Especially in LC-MS/MS lipidomic experiments this frequently leads to inaccurate or even incorrect annotations of lipid species. This problem is aggravated by a restricted selection of commercially available plasmanyl and plasmenyl standards. Importantly, automated analyses of LC-MS/MS datasets are particularly dependent on the databases used, which often prove to be incomplete with regard to ether lipids. Other analytical methods such as thin layer chromatography, on the other hand, are limited to measuring pools and ratios and cannot distinguish between individual molecular species ^12^, an important aspect that would be necessary to advance research in this field.

Here we demonstrate that the chemical properties of 1-O-alk-1’-enyl and 1-O-alkyl ether lipids generate a highly predictable chromatographic behavior of these compounds in reversed phase LC-MS/MS experiments that can be readily used for the univocal differentiation between plasmalogens and other ether lipid species. The exact characterization of these properties was facilitated by a comparative lipidomic analysis of plasmalogen-free PEDS knock-out mice with wild type littermate controls. Combined with the unique fragmentation properties of ether lipids, an exact molecular characterization of all relevant ether lipids could be carried out. This allowed us to map these lipids in detail across mouse tissues, thereby providing novel insights into their metabolism.

## Results

Highly reliable reference samples are required to validate the deconvolution of 1-O-alk-1’-enyl and 1-O-alkyl ether lipid features in LC-MS/MS experiments. As the selection of relevant commercial ether lipid and plasmalogen standards is still limited, we utilized mouse tissues as model system. The plasmalogen content in tissues of mice varies from 60 nmol/mg total protein in cerebellum and cerebrum to as little as 0.6-1.5 nmol/mg total protein in liver ^5^. Plasmalogen is depleted in all tissues of mice with absence of functional PEDS enzyme (Supplemental Figure 1). Thus, comparative analysis of ether lipid features between tissues of PEDS*-*deficient (ΔPEDS) mice and wild type (wt) controls represented the perfect model system to validate strategies for the comprehensive differentiation between plasmanyl and plasmenyl lipids. Here we first focused on kidney lipid extracts, as this tissue presented with the highest PEDS expression and activity levels ^5^.

Phospholipids were extracted and analyzed by LC-MS/MS as described previously ^13^ and the retention time and m/z characteristics of all features were compared. While some m/z overlap problems between plasmenyl-PC and phosphatidylserines or PE species can be readily resolved by measuring with mass resolutions larger than 21k and 14k, respectively, the truly isobaric nature of plasmanyl and plasmenyl PE/PC species remains a major analytical challenge. The similarity of these functionally distinct lipids in terms of mass to charge ratio and fragmentation behavior makes it especially difficult for automatic peak identification software to deliver correct results.

As illustrated in Figure 2A, wt kidney samples contained high levels of plasmenyl-PE species (left panel: PE(P-38:4), PE(P-38:5), PE(P-38:6) in red), which were absent in ΔPEDS kidneys, where instead the respective plasmanyl-PE species accumulated (right panel: PE(O-38:4), PE(O-38:5), PE(O-38:6) in blue). For other non-ether lipid species such as PE(36:2), PE(36:3), PE(36:4), and sphingomyelin(d34:1) we observed no differences in abundance as indicated by the black color in the overlay illustration (Figure 2A, center panel). For PC ether lipids a highly comparable behavior was observed (Figure 2B), however, with the striking difference that plasmanyl-PC species (PC(O-34:1), PC(O-34:2), PC(O-34:3)) were already much more abundant in the wt compared to plasmenyl-PC lipids (PC(P-34:1), PC(P-34:2), PC(P-34:3)). Such comparisons of ΔPEDS and wt tissues were further used to conduct the exact assignment of all alk-1’-enyl and alkyl lipids.

**Figure 2:**
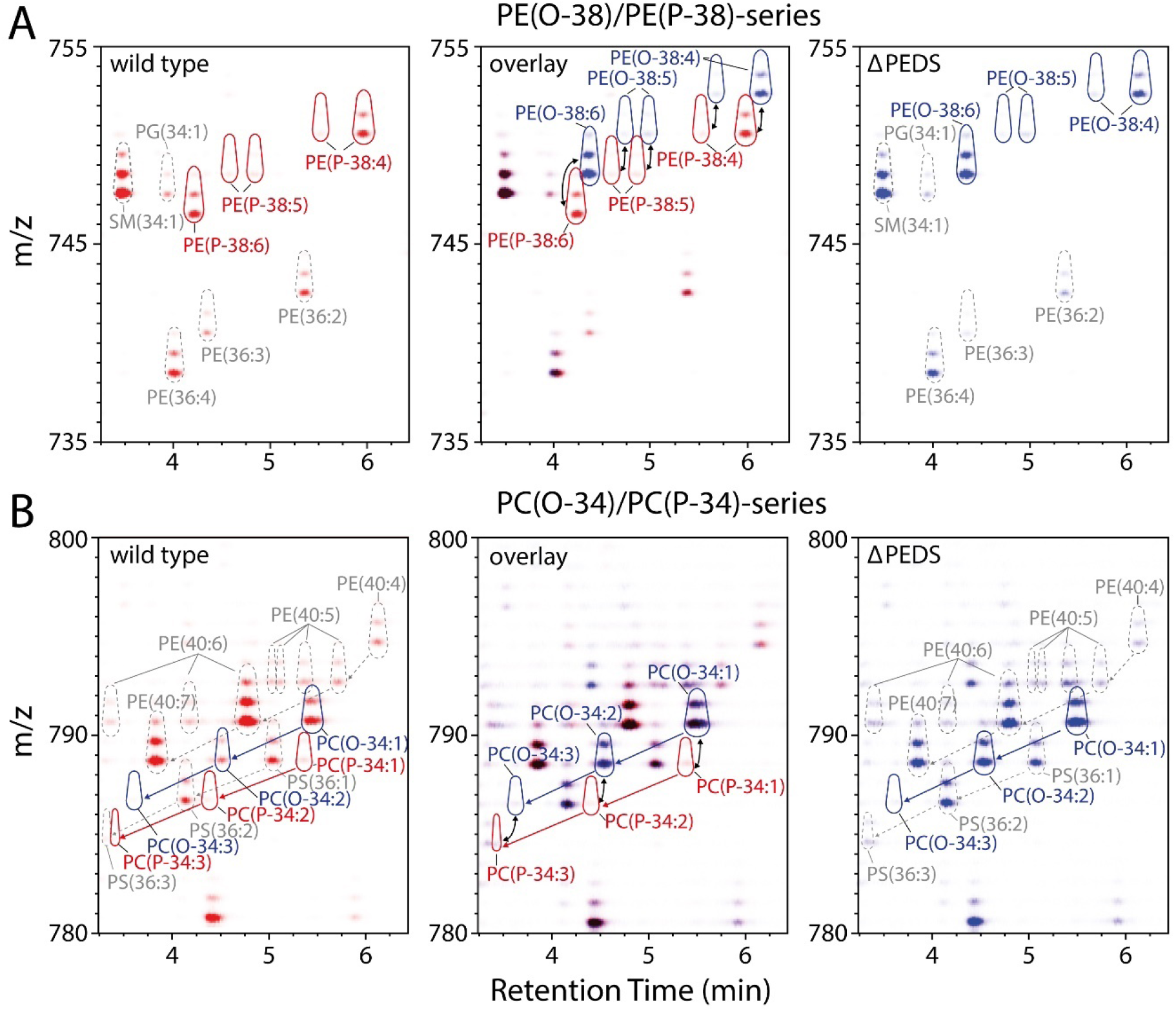
Annotation of plasmanyl and plasmenyl lipids by means of HPLC-MS/MS. A) 2D illustration of the MS1-data for wild type lipid extracts in the retention time range 3.25 - 6.5 min versus the mass window 735 – 755 m/z. This section mainly includes PE ether lipids with 38 and diacyl-PEs with 36 side chain carbons. Signals recorded in wild type and PEDS- deficient (ΔPEDS) kidneys are shown in red (left panel) and blue (right panel), respectively. Color intensity corresponds to the respective signal intensities as assigned by MZmine2. An overlay of wild type and ΔPEDS is shown in the central panel, with no differences being indicated in black. Black arrows indicate pairs of 1-O-alkyl and 1-O-alk-1’-enyl species that only differ in the vinyl ether double bond. Dashed black lines show retention time differences between selected features. Outline color scheme: red: 1-O-alk-1-enyl species; blue: 1-O-alkyl species; grey: diacyl species. B) Same as (A), but for the mass window 780 – 800 m/z. This section mainly includes PC ether lipids with 34 side chain carbons and diacyl-PEs with 40 side chain carbons. One representative example of at least three biological replicates is shown for A and B.

Next, we systematically analyzed the chromatographic properties of plasmalogens versus ether lipids and related these properties to the observed analytical behavior. In control kidney mainly 1-O-alk-1’-enyl species were detected (Figure 3A, red trace), while in ΔPEDS kidney 1-O-alkyl ether lipids dominated (Figure 3A, blue trace). At 750.6 *m/z* four isobaric ether lipid species were identified as [I] PE(O-18:1/20:4), [II] PE(P-18:0/20:4), [III] PE(O-16:0/22:5) and [IV] PE(P-16:0/22:4) (Figure 3A). This showed that the double bond at the Δ1 position of [II] and [IV] leads to a clearly different separation behavior than a double bond further back in the alkyl residue at the *sn*-1 [I] or in the acyl residue at the *sn-*2 position [III].

**Figure 3:**
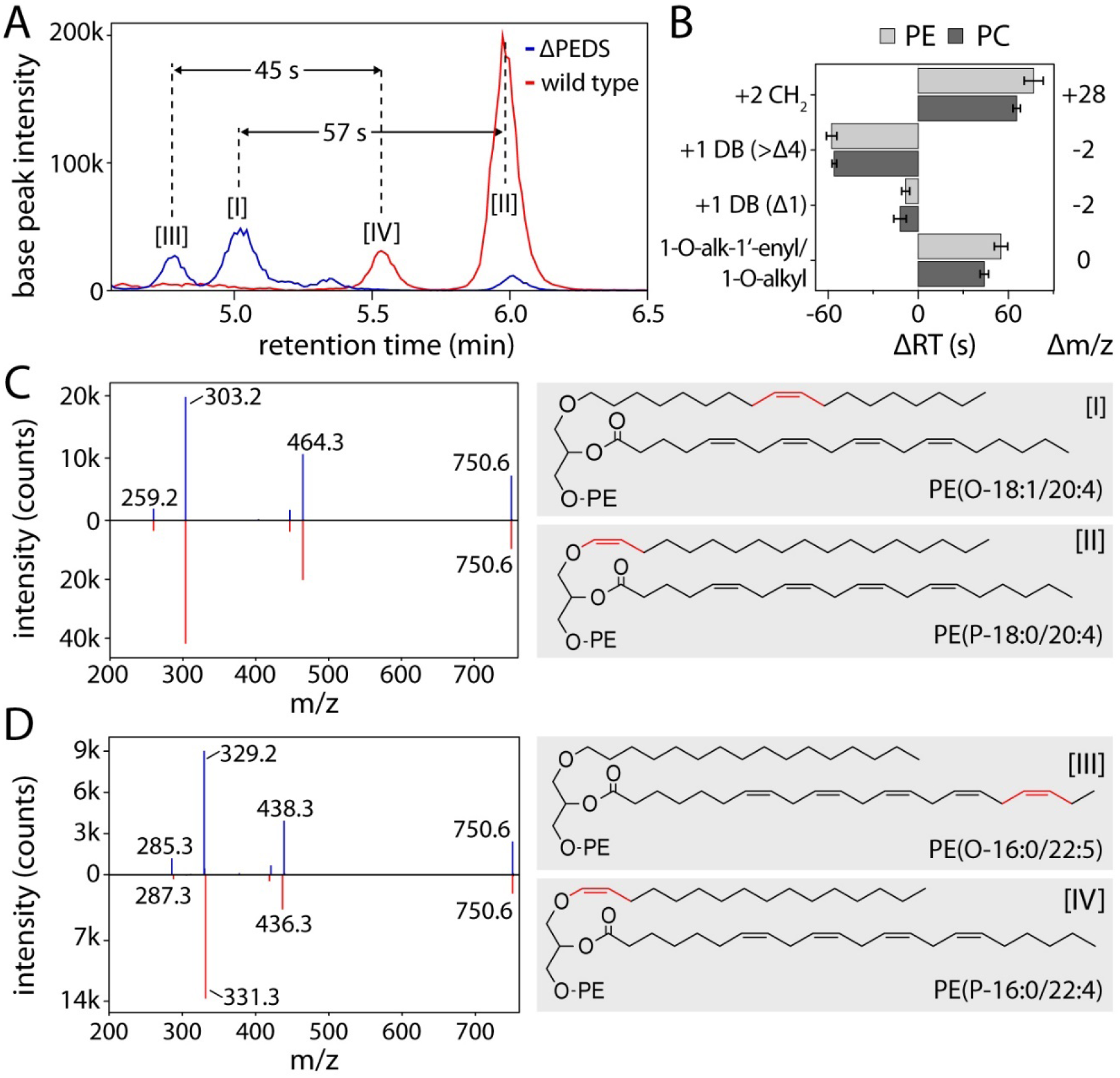
Discrimination between 1-O-alkyl and 1-O-alk-1’-enyl lipids in LC-MS/MS experiments. A) The isobaric ether lipid species [I] PE(O-18:1/20:4), [II] PE(P-18:0/20:4), [III] PE(O-16:0/22:5) and [IV] PE(P-16:0/22:4) could be separated by reversed phase chromatography, demonstrating that vinyl ether double bonds caused a retention time shift (ΔRT) different from double bonds further within the radyl residues. Blue trace: ΔPEDS, red trace wild type. B) Systematic analysis of ΔRT for acyl chain elongation (+2 CH_2_), an additional acyl chain double bond (+1 DB (>Δ4)), additional vinyl ether double bonds (+1 DB (Δ1)), and between 1-O-alk-1’-enyl and 1-O-alkyl lipids (1-O-alk-1’-enyl/1-O-alkyl) in 6-10 relevant lipid species each (n=12) allowed describing the impact of double bond positions and chain lengths on the elution time. C) Left panel: Collision-induced dissociation (CID) fragment spectra of [I] (blue) and [III] (red) are indistinguishable. One representative spectrum is shown each. Right panel: Structures of [I] and [III], with the differential double bond highlighted in red. D) Left panel: CID fragment spectra of [II] (blue) and [IV] (red) are distinguishable. One representative spectrum is shown each. Right panel: Structures of [II] and [IV], with the differential double bond highlighted in red.

This dataset was also used to characterize the general separation behavior of PE and PC lipids in the here used reversed phase chromatography system, by a pairwise comparison of 6-10 relevant lipid species (n=12) (Figure 3B, Supplemental Table 1). The elongation of a lipid side chain by two carbons resulted in a mass shift of +28 and a prolonged retention time of 71.1 ± 7.4 s. One additional double bond in an acyl or alkyl side chain caused a mass shift of −2 and a shortening of the retention time by 56.8 ± 2.8 s. However, an additional vinyl ether double bond did only reduce the retention time by 10.2 ± 3.9 s. In line with these observations alk-1’-enyl lipids elute 49.5 ± 6.7 s earlier than the corresponding isobaric alkyl species (45 s and 57 s in the examples of Figure 3A). This effect was observed in PE as well as PC and was thus largely lipid class independent.

Furthermore, our analysis revealed that a series of isobaric 1-O-alk-1’-enyl and 1-O-alkyl lipids such as (I) and (II) have identical fragmentation patterns in negative ESI mode (Figure 3C). Fragmentation behavior was as described in ^14^. Fragment spectra of PE-ether lipids in negative ESI mode generated *m/z* signals corresponding either to the neutral loss of the *sn*-2 acyl chain or the *sn*-2 residue itself. Common PC-ether lipid fragments were head group loss (−60 *m/z*) and low intensity *sn*-2 acyl chain signals. It follows that fragment spectra allow to assign double bonds to the *sn*-1 or *sn*-2 residues (329.2 *vs*. 331.3 *m/z* fragments, Figure 3D). However, an unequivocal discrimination into 1-O-alk-1’-enyl and 1-O-alkyl ether lipid can only be achieved by combining this information with the respective chromatographic behavior (Figure 3B).

In the next step, the obtained lipid species specific analytical behavior described above was utilized to comprehensively annotate and quantify plasmanyl and plasmenyl lipids in male and female mouse tissues. This analysis verified that in wt mice 1-O-alk-1’-enyl and 1-O-alkyl-species are predominantly found in the lipid classes of PE and PC. As shown in Figure 4A the profile of diacyl-PEs (left panel) is largely similar between male and female as well as ΔPEDS and wt kidneys. While 1-O-alk-1’-enyl-PEs (right panel) were abundant in the wt situation, they were largely absent in the ΔPEDS tissue, where we found 1-O-alkyl-PE species (center panel) being accumulated instead. A broad diversity of different 1-O-alk-1’-enyl-PE species could be identified in wt kidneys, but only a few of which made up most of the plasmalogen mass (i.e. PE(P-36:4), PE(P-38:4), PE(P-38:6), PE(P-40:6), and PE(P-40:7)). The same molecular lipid species, only lacking the vinyl ether bond, were accumulated in ΔPEDS kidneys. This shows a strong direct relationship between the availability of 1-O-alkyl precursors and 1-O-alk-1’-enyl products of the PEDS reaction.

**Figure 4:**
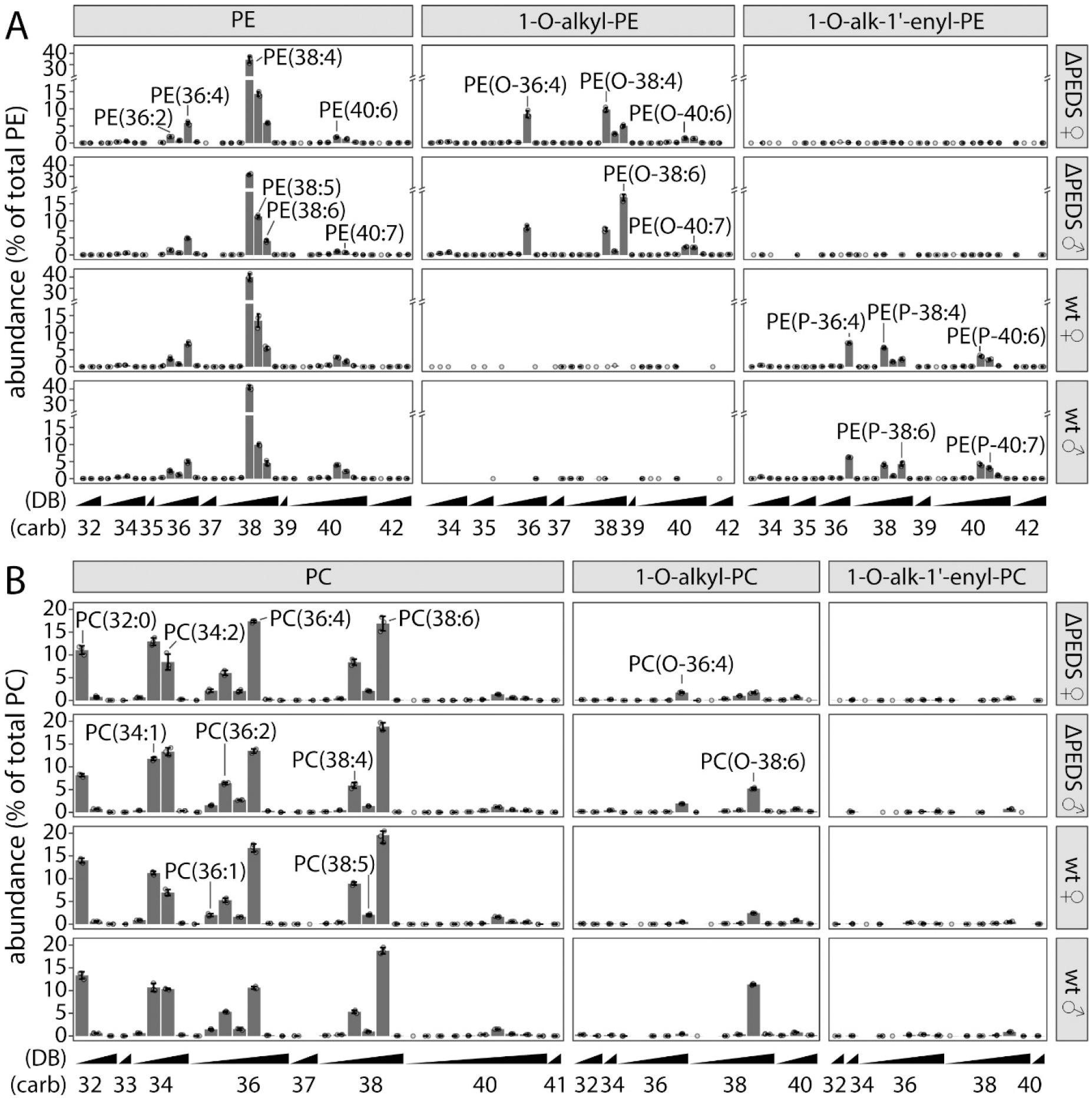
PE (A) and PC (B) in wild type and PEDS-deficient kidney. Lipid classes are subdivided into diacyl (left panel), 1-O-alkyl (center) and 1-O-alk-1’-enyl (right) species. Relative abundance profiles (in % of total PE or PC) are shown for wild type (wt) and PEDS-deficient (ΔPEDS) kidneys from female (♀) and male (♂) mice. Data is shown as mean ± SD (n=3). Selected lipid species are labeled: “O-” and “P-” indicating 1-O-alkyl and 1-O-alk-1’-enyl species, respectively. Numbers in parentheses give the total number of side chain carbon atoms (carb), followed by the number of double bonds (DB). Please note the axis break in (A).

Like diacyl-PE, also diacyl-PC profiles were largely unaffected by the knock-out of PEDS function (Figure 4B). However, in contrast to PEs, where plasmalogens dominated, we observed 13.5-times higher 1-O-alkyl-PC than 1-O-alk-1’-enyl-PC levels in wt kidneys. This is the quantified confirmation of the low wt levels of PC-plasmalogens as depicted in the example of Figure 4A. Additionally, 1-O-alk-1’-enyl-PC levels were further depleted in ΔPEDS samples. Importantly, the here described alterations of PE and PC ether lipid profiles were not restricted to the kidney, but could also be observed in cerebrum, heart and all other tested tissues (see Supplemental Dataset).

We used the here obtained lipidomics data for all tested tissues to describe overall plasmalogen and ether lipid status in wild type (wt) as well as PEDS-deficient (ΔPEDS) mouse tissues. Besides the above described kidney (KID), this includes cerebellum (CBL), cerebrum (CER), colon (COL), testis (TES), spleen (SPL), lung (LUN), heart (HRT), ovaries (OVA), skeletal muscle (SKM), and liver (LIV). Principal component analysis (PCA) of whole mouse lipid profiles (sequence of all tissue lipid profiles excluding TES and OVA) revealed that wt and ΔPEDS were separated by component 1 (Dim1) which captured 48.2% of the total variance (Figure 5A). Component 2 (Dim2) explained 17.5% of variance and distinguished between female (f) and male (m) mice. This indicates that even without reproductive organs enough differences remain that differentiate between the sexes. A further PCA was conducted between the lipid profiles of all individual tissues (Figure 5B). The main variance in this dataset was not explained by differences between female and male mice (left panel) but was mainly caused by differences in PEDS function (center panel) in combination with the nature of the respective tissue (right panel).

**Figure 5:**
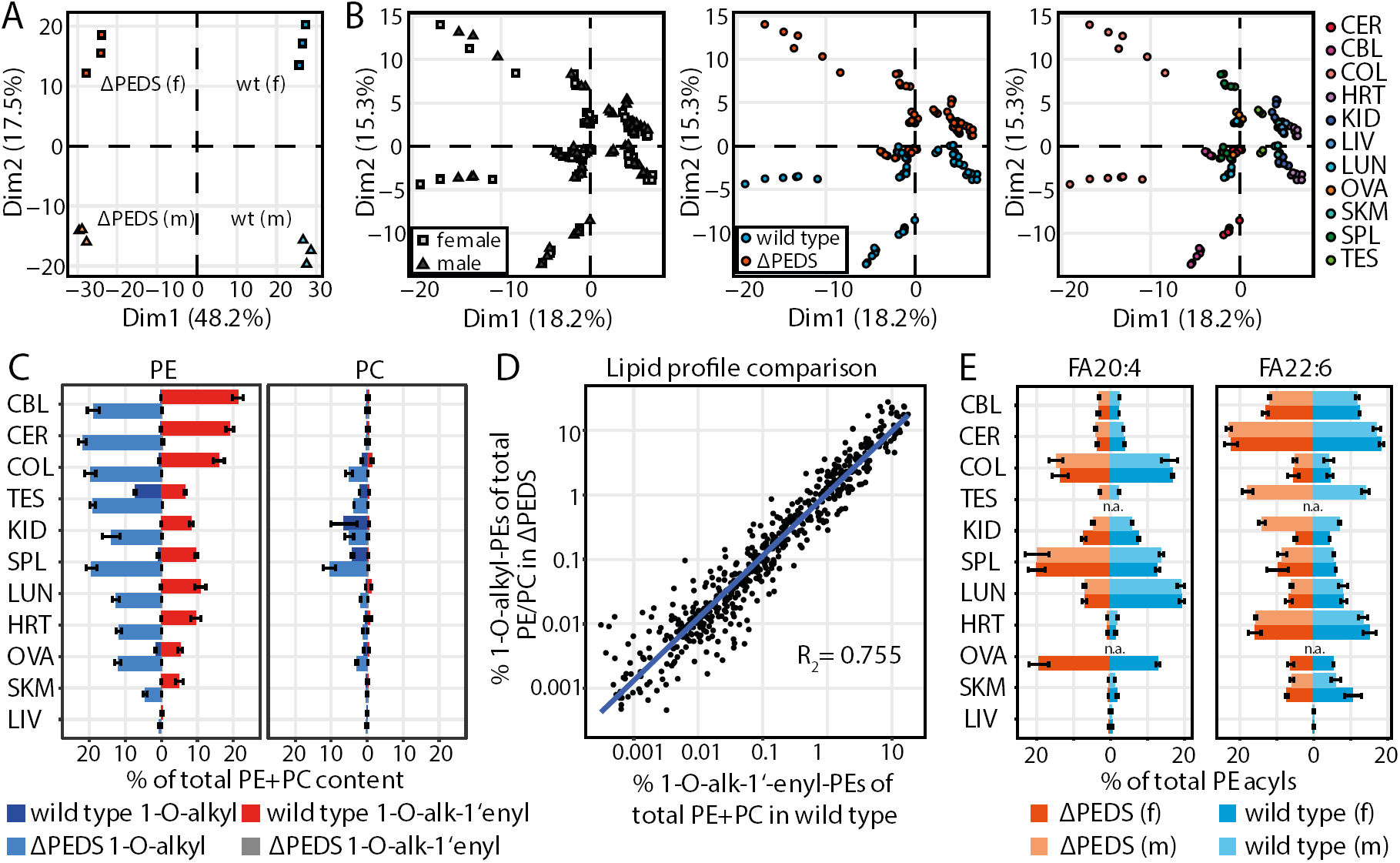
Ether lipid composition in wild type and PEDS-deficient mouse tissues. A) Principal component analysis (PCA) of a mouse-centered lipidomic dataset separates PEDS-deficient (ΔPEDS) and wild type mice in principal component 1 (Dim1; 48.2% of variance) and female and male mice in principal component 2 (Dim2; 48.2% of variance). B) PCA of individual tissue-centered dataset does not separate samples according to sexes (left), but discriminates between PEDS-deficient (ΔPEDS) and wild type tissues in principal component 2 (Dim2; 15.3% of variance; center). Additionaly the various tissues are separated by Dim2 and the principal component 1 (Dim1; 18.2% of variance; right). Tissues: CER: cerebrum, CBL: cerebellum; COL: colon; HRT: heart; KID: kidney; LIV: liver; LUN: lung; OVA: ovaries; SKM: skeletal muscle; SPL: spleen; TES: testis. C) Content of ether lipids in wild type and ΔPEDS tissues. Percentage of 1-O-alk-1’-enyl (red: wild type; orange: ΔPEDS) and 1-O-alkyl (blue: wild type; light blue: ΔPEDS) species related to total PE and PC lipids. Data shown as mean +/- SD (n=3). D) Correlation between 1-O-alk-1’-enyl species in wild type and their corresponding 1-O-alkyl species in ΔPEDS. Each dot represents one 1-O-alk-1’-enyl - 1-O-alkyl pair in a tissue. Data includes all tissues shown in (B). E) Content of fatty acyls (FA) 20:4 (left panel) an FA 22:6 (right panel), relative to the respective total PE acyl pool (excluding 1-O-akyl and 1-O-alk-1-enyl residues), shown for all tissues listed in (B) ordered according to (C). Coloring scheme: red: PEDS-deficient female; light red: PEDS-deficient male; blue: wild type female; light blue: wild type male. Data shown as mean ± SD (n=3).

Next, we analyzed the total 1-O-alkyl and 1-O-alk-1’-enyl content in tissues. In wild type mice, cerebrum and cerebellum had the highest total ether lipid content (22.1% and 19.9% of total PC and PE, respectively), which to a large extent consisted of 1-O-alk-1’-enyl-PEs (Figure 5C, red bars). In contrast, liver was almost depleted of ether lipids (0.7% of total PC and PE). PE ether lipids were notably more abundant than PC ether lipids across all tissues. Typically, only low levels of 1-O-alkyl-PE species were quantified, with the exception of testis and ovaries. Increased levels of PC ether lipids were present as 1-O-alkyl species in kidney, spleen and colon (Figure 5C, blue bars). Upon PEDS-deficiency we observed a quantitative depletion of 1-O-alk-1’-enyl species (Figure 5C, grey bars). Instead, 1-O-alkyl-PEs accumulated in ΔPEDS tissues (Figure 5C, light blue bars), partially to even higher levels than their 1-O-alk-1’-enyl counterparts in wt, as for example in spleen (2-fold), testis (2.9-fold), and ovaries (2.3-fold). A comparable effect, only at a significantly lower basic level, could also be measured with PCs. When comparing the lipid species profiles of wild type 1-O-alk-1’-enyl-PEs with the respective 1-O-alkyl-PEs in ΔPEDS we observed a consistent correlation between the molecular structures of these ether lipids in all tissues (Figure 5D). This showed that the profile similarity observed in kidney (Figure 4) follows a general, tissue independent behavior.

The fragmentation behavior of PE lipids and a high MS/MS coverage allowed to carry out a radyl chain specific analysis in this lipid class. Side chain information for PEs was automatically extracted from 25,923 relevant fragment spectra and used to determine the overall acyl composition in diacyl-PEs, and the *sn*-1/*sn*-2 specific substitution patterns in ether lipids. The *sn*-1 position of ether lipids was substituted mainly by 16:0, 18:0, and 18:1 fatty alcohols (Supplemental Dataset). In contrast, the major *sn*-2 residues in plasmanyl- and plasmenyl-PEs were fatty acyls (FA) FA 20:4 and FA 22:6, which was in line with the described preference for arachidonic and docosahexaenoic acid of these lipids (Supplemental Dataset). Their distribution was, however, not homogenous across tissues. While cerebrum, cerebellum, and heart were characterized by a high FA 22:6 and a low FA 20:4 levels, we observed the opposite in colon and spleen (Figure 5E). Also, some of the gender-specific differences could again be observed in this analysis. The content of FA 22:6 was considerably higher in male kidneys than in female, whereas FA 20:4 was more common in female kidneys than in male (Figure 5E). This is in line with the differences we observed for PE(O-38:4), PE(O-38:6), PE(P-38:6), and PE(P-38:6) in the lipid composition of male and female kidneys (compare Figure 4).

## Discussion

We report a reliable strategy for discriminating between plasmanyl and plasmenyl ether lipids in LC-MS/MS workflows. By exploiting the fully predictable changes in the chromatographic properties of these lipids, their behavior can be extrapolated from just a few reference features. Importantly, this approach does not require specialized mass spectrometric infrastructure and can in principle also be carried out on low resolution instruments without fragmentation capacity. The consistent implementation of these principles can contribute strongly towards advancing with the core goals of comprehensive lipidomics, which are to achieve highest possible coverage in terms of identification and quantification coupled with complete structural elucidation. Such an exact assignment of ether lipid species, despite their natural abundance, is only rarely carried out in lipidomics experiments.

A reliable discrimination between 1-O-alkyl and 1-O-alk-1’-enyl lipids is essential for phospholipid research. Ether lipids make up a considerable portion of the phospholipid mass in mammals as shown by others ^2,1^ and validated here (Figure 5B). Furthermore, ether lipid deficiency is thought to be the main cause of the phenotype of rhizomelic chondrodysplasia punctata and the respective features of Zellweger spectrum disorders ^15,3^. Still, the functional role of the vinyl ether double bond remains elusive ^16^. Also, it has not been studied in detail whether plasmalogen depletion – reported in some neurodegenerative diseases ^6^ – is accompanied by changes in 1-O-alkyl lipids in a parallel or opposing manner ^16^. The comprehensive analytical solution for studying ether lipids should enable more detailed exploration of the functions of this lipid class.

A major bottleneck is represented by the small number of commercially available ether lipid standards. For example, no pair of 1-O-alkyl and 1-O-alk-1’-enyl lipids with an otherwise similar side chain and head group configuration is currently available from Avanti Polar Lipids (Alabaster, USA, May 2020), with 1-O-alkyl-PE lipids particularly under-represented. However, in order to systematically elucidate the analytical properties of molecular 1-O-alkyl and 1-O-alk-1’-enyl ether lipid species reliable sample material is needed. Due to the recent identification of the *PEDS* gene that introduces the vinyl ether double bond and thus forms plasmalogens ^4,5^, we have been able to generate plasmalogen-free and 1-O-alkyl ether lipid enriched lipid extracts. This made extensive validation of ether lipid annotation in LC-MS/MS experiments possible.

As demonstrated in Figure 2 and Figure 3, 1-O-alkyl and 1-O-alk-1’-enyl lipids cannot be readily discriminated purely on the basis of exact mass and MS/MS fragment spectra. While ESI in positive mode only provides strongly limited side chain information, the here used negative mode gives specific details about the *sn*-1 and *sn*-2 residues. This is because in contrast to acyl chains, 1-O-alkyls and 1-O-alk-1’-enyls are exceptionally resistant against fragmentation. In some cases, if no double bond is found at the *sn-*1 position at all, this can directly point towards the identity of the ether lipid. In general, however, further information is required in order to differentiate between a vinyl ether at the Δ1 position or a double bond further within the fatty acyl chain. Using a reversed phase LC-gradient, we here demonstrate that a vinyl ether-specific retention time offset can be utilized for this differentiation (Figure 3). While other double bond positions lead to a mutually similar change in the retention time, isobaric vinyl ethers can be readily separated by approx. 50 s, even in short reversed phase gradients. This behavior was consistent for different molecular lipid species as well as lipid classes and is therefore predictable, which makes it immediately relevant for automated analyses of LC-MS/MS datasets. The benefits of this principle are demonstrated by its successful application in the lipidomics analysis of various mouse tissues (Figure 5).

It can be speculated that the same physiological properties that functionally differentiate plasmalogens from other ether lipids – e.g. with regard to their impact on membrane fluidity ^17^ – are the reason for the different chromatographic behavior of these lipids. These migration properties were already exploited in 2D-thin layer chromatography experiments ^18^ but did not allow separation of 1-O-alkyl from diacyls lipids without the use of chemical degradation procedures. Similarly, high-resolution ^31^P nuclear magnetic resonance cannot differentiate between 1-O-alkyl from diacyl lipids although it recognizes plasmalogens as a separate feature ^19^. The advantage of attributing for the systematic behavior shown here is that the laborious individual confirmation of ether lipid species (for example using hydrochloric acid hydrolyzed reference material) which is carried out in some particularly well-controlled studies ^20^ is significantly simplified.

In many academic and commercial lipidomic experiments the annotation of ether lipid species is based on assumptions. This problem is not only restricted to the differentiation between 1-O-alkyl and 1-O-alk-1’-enyl species but also extends, for example, to the exact assignment of the *sn*-1 and *sn*-2 side chains or double bond positions ^21^. Although desirable, it is rare that such ambiguities are communicated precisely and are considered in further steps of data analysis ^10^. This leads to the situation that even in otherwise well performed systematic lipidomic studies the abundant class of ether lipids is not mentioned ^22^. Even in the absence of supporting structural data the identification as a plasmalogen is typically given priority over other ether lipids. Fortunately, this default preference is in agreement with the strong overrepresentation of 1-O-alk-1’-enyl-PE species as found in our mouse tissue analysis (Figure 5) and elsewhere ^20^. In contrast, for PC lipids we observe a considerable proportion of 1-O-alkyl species across tissues (Figure 5). Special caution is also required when studying conditions that cause a disturbed plasmanyl and plasmenyl metabolism. Such conditions include for example oxidative stress, for which vinyl ether double bonds are especially susceptible ^23,24^, or the here presented extreme case of PEDS-deficiency.

The fraction of PE and PC ether lipids in respect to total biomass or total phospholipid content has long been of scientific interest and has been studied with special focus on 1-O-alk-1’-enyl species in various organisms and tissues ^2,25^. Several of these studies report conflicting results. For example, the proportion of PE plasmalogens in mammalian heart tissue has been reported to range from 12% to 48% of all phospholipids, with a trend towards higher values for larger organisms ^25^. Particularly high PC plasmalogen levels have been described in human heart tissue ^1515^, in some cases even higher than their PE counterparts ^26^. This led to the conclusion that 1-O-alk-1’-enyl PEs are typically about one order of magnitude more abundant than 1-O-alk-1’-enyl PCs, with the exception of muscle and heart ^27^. In contrast, in the current study we observed a PE to PC plasmalogen ratio of 11.5 in normal murine heart tissue. Many early studies focusing on ether lipids relied on methods such as thin layer chromatography or selective plasmalogen cleavage and derivatization ^28^. The presence of free fatty aldehydes has the potential to distorted the results of such experiments and zo miscalculate plasmalogen levels ^29^.

Indeed, in a more recent LC-MS/MS based study a PE to PC plasmalogen ratio of about 20 was recorded in hearts from eight-month-old rats ^20^. However, not all differences can and should be attributed to technical inaccuracies and differences. It has been shown that tissue ether lipid levels – including heart – were highly responsive to the dietary alkyl-glycerol content in wild type as well as ether lipid deficient mice ^30^. Thus, it is to be expected that a strong nutritional component exists in addition to the species- and tissue-dependent differences. Much of this is still unexplored and requires precise, robust and reliable analytical strategies for future studies.

While the total amount of plasmalogens can be readily measured by cleavage of the vinyl ether bond in hydrochloric acid and derivatization of the resulting aldehyde to a dimethyl acetal and MS detection ^30^, or derivatization a fluorescent hydrazone and fluorescence detection ^29^, 1-O-alkyl lipid concentrations are monitored less frequently, although they may be equally important. As is demonstrated in Figure 5C, PE ether lipids occur mostly as plasmalogens except for testis where we find equal amounts of 1-O-alkyl and 1-O-alk-1’-enyl PE species. In testis, seminolipid is formed which is a complex ether lipid without a vinyl ether bond and is required for male fertility and sperm maturation ^31^. Remarkably, male PEDS knock-out mice are fertile ^32,5^ underlining the notion that no vinyl ether bond is required for seminolipid function. In PEDS-deficient tissues such as spleen, testis, and ovaries 1-O-alkyl lipids accumulate to even higher levels than the total ether lipid content in the wild type (Figure 5B). This has to be accounted for when interpreting respective phenotypes. Since many 1-O-alkyl species only accumulate under specific conditions, it is particularly important that the analytical behavior shown here (Figure 3) can be utilized to find and identify them.

Our work adds a promising and easy-to-implement strategy to map all types of ether lipids and understand their role in physiology and pathophysiology in the future, by allowing a more comprehensive characterization of their relative and absolute quantities and studying their regulation in healthy and diseased states.

## Methods

### Breeding of *Tmem189*-deficient mice and harvest of mouse tissues

Mice deficient in the PEDS enzyme (Tmem189tm1a(KOMP)Wtsi mice, Wellcome Sanger Institute, Hinxton, Cambridge, UK) were bred according to (Werner et al. 2020) and was approved by the Austrian Federal Ministry of Education, Science and Research (BMBWF-66.011/0100-V/3b/2019). Mice were housed in individual ventilated cages with nesting material, in a 12 h/12 h light/dark cycle with standard chow (Sniff Spezialdiäten GmbH, standard chow V1534-300, autoclaved at site) and water ad libitum. They were maintained on C57bl/6N genetic background and genotyped using primers Tmem189_35635F (GCGTGTCCTGCTGAGACTTG) and CAS_R1_Term (TCGTGGTATCGTTATGCGCC) for the transgenic allele and Tmem189_35635F and Tmem189-35635R (CATCCCACCTATCCCACCTG) for the wildtype allele. Tissues were harvested at 8 weeks of age from female and male homozygous *Peds*-deficient mice and their wild type littermates after sacrificing the animals by cervical dislocation. Tissues were snap frozen in liquid nitrogen and stored at −80 °C until further analysis.

### Sample extraction and preparation

Sample preparation was performed as previously described ^29^. Briefly tissues were homogenized in PBS with an Ultra-Turrax (T10, IKA, Staufen, Germany). The protein content was then measured with a standard Bradford assay depending on a BSA serial dilution for quantification. For lipid extraction per tissue 500 µg total protein aliquots were extracted following the Folch method. Dried lipid extracts were finally dissolved in 100 µl 1/1 (v/v) acetonitrile/ethanol and stored at −20 °C. Aliquots with lipids respective to 200 µg (and for brain tissues 150 µg) protein content were dried under N_2_-flow, dissolved in 100 µl HPLC starting conditions and subsequently measured.

### LC-MS/MS analysis

The PC/PE analysis was performed as described in ^33^, with the modifications described in ^13^. 10 µl of the samples were injected by a Dionex Ultimate 3000 HPLC (Thermofisher Scientific Inc, Waltham, USA) with a flowrate of 0.4 ml/min and a column oven temperature of 50 °C. The HPLC was operated in reversed phase mode using an Agilent Poroshell 120 EC-C8 2.7mm 2.1×100mm column (Agilent Technologies, Santa Clara, USA). For gradient elution running Solvent A (4/6 H_2_O/ammonium format, 0.2% formic acid) and running Solvent B (9/1 isopropanol/acetonitrile, 10 mM ammonium formate, 0.2% formic acid) were used. First, from 0 to 2 minutes, an isocratic flow with 54% Solvent A was applied, followed by a linear decrease from 54% to 28% Solvent A from minute 2 to minute 22. Afterwards the column was washed and again equilibrated at a starting condition of 54% Solvent A. Further, lipids were ionized through negative electrospray ionization (275 °C capillary temperature, with mass range of 460-1650 m/z) in a LTQ Velos MS (Thermo Fisher Scientific Inc., Waltham, USA), whereas MS2 settings were variable depending on the expected PL. For general LC-MS parameters see Supplemental Table 2 and 3.

### Data analysis

Raw data visualization was done in MZmine 2 version 2.53 ^34^, for all further analysis steps we utilized our in-house data extraction pipeline written in R ^35–37^. Features were extracted in a targeted manner according to a retention time adjusted manually created peak list template (see Supplemental Table 4). Peak identities were confirmed by MS2 fragmentation spectra, retention time, m/z ratio, and cross-validation with PL lipid standards. The pipeline considers the base - and first isotope peaks for quantification. For boundary definition of the peaks we assumed gaussian typologies. These peaks were afterwards baseline corrected, filtered, and quantified by an external standard series. Peaks not fulfilling user defined criteria were rejected and overlapping peaks attributed to the same molecular species were resolved in an automated manner, while overlaps belonging to different isobaric lipid species were highlighted and manually resolved by consideration of MS2 data, retention time, and comparison between sample groups. To account for sample-sample variations lipids were normalized to total analyzed lipid content per sample (PE+PC) and used for data visualisation.

### Data availability

Datasets have been deposited (see Supplemental Dataset).

## Acknowledgements

The authors thank Petra Loitzl and Nina Madl (Institute of Biological Chemistry) for expert technical help. This work was supported by a Medical University of Innsbruck, Austria Start grant to M.A.K., the Austrian Science Fund, Austria (FWF) projects P29551 to E.R.W and P30800 to K.W.

## Author contributions

J.K., E.R.W., J.Z., K.W., and M.A.K. designed the study; J.K., S.S., and K.L. performed experimental work; J.K., M.A.K. analyzed data; M.A.K., E.R.W., J.K., K.W., and Y.W. wrote the paper with input from all authors.

## References

(1) Nagan, N.; Zoeller, R. A., Plasmalogens: biosynthesis and functions, Progress in lipid research, 2001, DOI: 10.1016/s0163-7827(01)00003-0.

(2) Braverman, N. E.; Moser, A. B., Functions of plasmalogen lipids in health and disease, Biochimica et biophysica acta, 2012, DOI: 10.1016/j.bbadis.2012.05.008.

(3) Waterham, H. R.; Ferdinandusse, S.; Wanders, R. J. A., Human disorders of peroxisome metabolism and biogenesis, Biochimica et biophysica acta, 2016, DOI: 10.1016/j.bbamcr.2015.11.015.

(4) Gallego-García, A.; Monera-Girona, A. J.; Pajares-Martínez, E.; Bastida-Martínez, E.; Pérez-Castaño, R.; Iniesta, A. A.; Fontes, M.; Padmanabhan, S.; Elías-Arnanz, M., A bacterial light response reveals an orphan desaturase for human plasmalogen synthesis, Science (New York, N.Y.), 2019, DOI: 10.1126/science.aay1436.

(5) Werner, E. R.; Keller, M. A.; Sailer, S.; Lackner, K.; Koch, J.; Hermann, M.; Coassin, S.; Golderer, G.; Werner-Felmayer, G.; Zoeller, R. A.; Hulo, N.; Berger, J.; Watschinger, K., The TMEM189 gene encodes plasmanylethanolamine desaturase which introduces the characteristic vinyl ether double bond into plasmalogens, Proceedings of the National Academy of Sciences of the United States of America, 2020, DOI: 10.1073/pnas.1917461117.

(6) Han, X.; Holtzman, D. M.; McKeel, D. W., Plasmalogen deficiency in early Alzheimer’s disease subjects and in animal models: molecular characterization using electrospray ionization mass spectrometry, Journal of neurochemistry, 2001, DOI: 10.1046/j.1471-4159.2001.00332.x.

(7) Keller, M. A.; Zander, U.; Fuchs, J. E.; Kreutz, C.; Watschinger, K.; Mueller, T.; Golderer, G.; Liedl, K. R.; Ralser, M.; Kräutler, B.; Werner, E. R.; Marquez, J. A., A gatekeeper helix determines the substrate specificity of Sjögren-Larsson Syndrome enzyme fatty aldehyde dehydrogenase, Nature communications, 2014, DOI: 10.1038/ncomms5439.

(8) Watschinger, K.; Keller, M. A.; Golderer, G.; Hermann, M.; Maglione, M.; Sarg, B.; Lindner, H. H.; Hermetter, A.; Werner-Felmayer, G.; Konrat, R.; Hulo, N.; Werner, E. R., Identification of the gene encoding alkylglycerol monooxygenase defines a third class of tetrahydrobiopterin-dependent enzymes, Proceedings of the National Academy of Sciences of the United States of America, 2010, DOI: 10.1073/pnas.1002404107.

(9) Wu, L.-C.; Pfeiffer, D. R.; Calhoon, E. A.; Madiai, F.; Marcucci, G.; Liu, S.; Jurkowitz, M. S., Purification, identification, and cloning of lysoplasmalogenase, the enzyme that catalyzes hydrolysis of the vinyl ether bond of lysoplasmalogen, J. Biol. Chem., 2011, DOI: 10.1074/jbc.M111.247163.

(10) Holčapek, M.; Liebisch, G.; Ekroos, K., Lipidomic Analysis, Analytical chemistry, 2018, DOI: 10.1021/acs.analchem.7b05395.

(11) Liebisch, G.; Ekroos, K.; Hermansson, M.; Ejsing, C. S., Reporting of lipidomics data should be standardized, Biochimica et biophysica acta. Molecular and cell biology of lipids, 2017, DOI: 10.1016/j.bbalip.2017.02.013.

(12) Owens, K., A two-dimensional thin-layer chromatographic procedure for the estimation of plasmalogens, The Biochemical journal, 1966, DOI: 10.1042/bj1000354.

(13) Oemer, G.; Koch, J.; Wohlfarter, Y.; Alam, M. T.; Lackner, K.; Sailer, S.; Neumann, L.; Lindner, H. H.; Watschinger, K.; Haltmeier, M.; Werner, E. R.; Zschocke, J.; Keller, M. A., Phospholipid Acyl Chain Diversity Controls the Tissue-Specific Assembly of Mitochondrial Cardiolipins, Cell reports, 2020, DOI: 10.1016/j.celrep.2020.02.115.

(14) Hsu, F.-F.; Turk, J., Differentiation of 1-O-alk-1’-enyl-2-acyl and 1-O-alkyl-2-acyl glycerophospholipids by multiple-stage linear ion-trap mass spectrometry with electrospray ionization, Journal of the American Society for Mass Spectrometry, 2007, DOI: 10.1016/j.jasms.2007.08.019.

(15) Heymans, H. S.; Schutgens, R. B.; Tan, R.; van den Bosch, H.; Borst, P., Severe plasmalogen deficiency in tissues of infants without peroxisomes (Zellweger syndrome), Nature, 1983, DOI: 10.1038/306069a0.

(16) Dorninger, F.; Forss-Petter, S.; Berger, J., From peroxisomal disorders to common neurodegenerative diseases - the role of ether phospholipids in the nervous system, FEBS letters, 2017, DOI: 10.1002/1873-3468.12788.

(17) Koivuniemi, A., The biophysical properties of plasmalogens originating from their unique molecular architecture, FEBS letters, 2017, DOI: 10.1002/1873-3468.12754.

(18) Zoeller, R. A.; Grazia, T. J.; LaCamera, P.; Park, J.; Gaposchkin, D. P.; Farber, H. W., Increasing plasmalogen levels protects human endothelial cells during hypoxia, American journal of physiology. Heart and circulatory physiology, 2002, DOI: 10.1152/ajpheart.00524.2001.

(19) Kimura, T.; Kimura, A. K.; Ren, M.; Berno, B.; Xu, Y.; Schlame, M.; Epand, R. M., Substantial Decrease in Plasmalogen in the Heart Associated with Tafazzin Deficiency, Biochemistry, 2018, DOI: 10.1021/acs.biochem.8b00042.

(20) Pradas, I.; Huynh, K.; Cabré, R.; Ayala, V.; Meikle, P. J.; Jové, M.; Pamplona, R., Lipidomics Reveals a Tissue-Specific Fingerprint, Frontiers in physiology, 2018, DOI: 10.3389/fphys.2018.01165.

(21) Rustam, Y. H.; Reid, G. E., Analytical Challenges and Recent Advances in Mass Spectrometry Based Lipidomics, Analytical chemistry, 2018, DOI: 10.1021/acs.analchem.7b04836.

(22) Jain, M.; Ngoy, S.; Sheth, S. A.; Swanson, R. A.; Rhee, E. P.; Liao, R.; Clish, C. B.; Mootha, V. K.; Nilsson, R., A systematic survey of lipids across mouse tissues, American journal of physiology. Endocrinology and metabolism, 2014, DOI: 10.1152/ajpendo.00371.2013.

(23) Jansen, G. A.; Wanders, R. J., Plasmalogens and oxidative stress: evidence against a major role of plasmalogens in protection against the superoxide anion radical, Journal of inherited metabolic disease, 1997, DOI: 10.1023/A:1005321910248.

(24) Lessig, J.; Fuchs, B., Plasmalogens in biological systems: their role in oxidative processes in biological membranes, their contribution to pathological processes and aging and plasmalogen analysis, Current medicinal chemistry, 2009, DOI: 10.2174/092986709788682164.

(25) Snyder, F.; Baumann, W. J., Ether lipids: chemistry and biology; Academic Press: New York, 1972.

(26) Panganamala, R. V.; Horrocks, L. A.; Geer, J. C.; Cornwell, D. G., Positions of double bonds in the monounsaturated alk-1-enyl groups from the plasmalogens of human heart and brain, Chemistry and Physics of Lipids, 1971, DOI: 10.1016/0009-3084(71)90031-4.

(27) Farooqui, A. A.; Horrocks, L. A., Plasmalogens: workhorse lipids of membranes in normal and injured neurons and glia, The Neuroscientist : a review journal bringing neurobiology, neurology and psychiatry, 2001, DOI: 10.1177/107385840100700308.

(28) Gottfried, E. L.; Rapport, M. M., The Biochemistry of Plasmalogens: I. ISOLATION AND CHARACTERIZATION OF PHOSPHATIDAL CHOLINE, A PURE NATIVE PLASMALOGEN, J. Biol. Chem., 1962, 237 (2), 329–333.

(29) Werner, E. R.; Keller, M. A.; Sailer, S.; Seppi, D.; Golderer, G.; Werner-Felmayer, G.; Zoeller, R. A.; Watschinger, K., A novel assay for the introduction of the vinyl ether double bond into plasmalogens using pyrene-labeled substrates, Journal of lipid research, 2018, DOI: 10.1194/jlr.D080283.

(30) Brites, P.; Ferreira, A. S.; da Silva, T. F.; Sousa, V. F.; Malheiro, A. R.; Duran, M.; Waterham, H. R.; Baes, M.; Wanders, R. J. A., Alkyl-glycerol rescues plasmalogen levels and pathology of ether-phospholipid deficient mice, PloS one, 2011, DOI: 10.1371/journal.pone.0028539.

(31) Tanphaichitr, N.; Kongmanas, K.; Faull, K. F.; Whitelegge, J.; Compostella, F.; Goto-Inoue, N.; Linton, J.-J.; Doyle, B.; Oko, R.; Xu, H.; Panza, L.; Saewu, A., Properties, metabolism and roles of sulfogalactosylglycerolipid in male reproduction, Progress in lipid research, 2018, DOI: 10.1016/j.plipres.2018.08.002.

(32) Ingham, N. J.; Pearson, S. A.; Vancollie, V. E.; Rook, V.; Lewis, M. A.; Chen, J.; Buniello, A.; Martelletti, E.; Preite, L.; Lam, C. C.; Weiss, F. D.; Powis, Z.; Suwannarat, P.; Lelliott, C. J.; Dawson, S. J.; White, J. K.; Steel, K. P., Mouse screen reveals multiple new genes underlying mouse and human hearing loss, PLoS biology, 2019, DOI: 10.1371/journal.pbio.3000194.

(33) Oemer, G.; Lackner, K.; Muigg, K.; Krumschnabel, G.; Watschinger, K.; Sailer, S.; Lindner, H.; Gnaiger, E.; Wortmann, S. B.; Werner, E. R.; Zschocke, J.; Keller, M. A., Molecular structural diversity of mitochondrial cardiolipins, Proceedings of the National Academy of Sciences of the United States of America, 2018, DOI: 10.1073/pnas.1719407115.

(34) Pluskal, T.; Castillo, S.; Villar-Briones, A.; Oresic, M., MZmine 2: modular framework for processing, visualizing, and analyzing mass spectrometry-based molecular profile data, BMC bioinformatics, 2010, DOI: 10.1186/1471-2105-11-395.

(35) FBernd ischer, S. N., mzR; Bioconductor, 2017, DOI: 10.18129/B9.bioc.mzR.

(36) R Core Team, R: A language and environment for statistical computing; R Foundation for Statistical Computing: Vienna, Austria, 2020.

(37) Wickham, H.; Averick, M.; Bryan, J.; Chang, W.; McGowan, L.; François, R.; Grolemund, G.; Hayes, A.; Henry, L.; Hester, J.; Kuhn, M.; Pedersen, T.; Miller, E.; Bache, S.; Müller, K.; Ooms, J.; Robinson, D.; Seidel, D.; Spinu, V.; Takahashi, K.; Vaughan, D.; Wilke, C.; Woo, K.; Yutani, H., Welcome to the Tidyverse, JOSS, 2019, DOI: 10.21105/joss.01686.

(38) Liebisch, G.; Vizcaíno, J. A.; Köfeler, H.; Trötzmüller, M.; Griffiths, W. J.; Schmitz, G.; Spener, F.; Wakelam, M. J. O., Shorthand notation for lipid structures derived from mass spectrometry, Journal of lipid research, 2013, DOI: 10.1194/jlr.M033506.

