## Supplementary Material for "Unequivocal mapping of molecular ether lipid species by LC-MS/MS in plasmalogen-deficient mice"

***Supplemental Dataset: Contains additional information and data for all 11 different mouse tissues analyzed.*** *For all tissues you will find there a PC and PE lipid profiles plot, a radyl and acyl chain plot for PE lipid species, and the data used for most of the analyses presented in this work. See Mendeley Data at http://dx.doi.org/10.17632/r5twyf6bnn.1*

***Supplemental Table 1: Table detailing the systematic analysis of retention time differences (ΔRT).*** *We analyzeda ΔRTs for acyl chain elongation (+2 CH2), an additional acyl chain double bond (+1 DB (>Δ4)), additional vinyl ether double bonds (+1 DB (Δ1)), and between 1-O-alk-1'-enyl and 1-O-alkyl lipids in 6-10 relevant lipid species, which allowed us to number the impact of double bond positions and chain lengths on lipid retardation. ΔRT in seconds was calculated as* $\Delta RT\ [\mathrm{s}] = rt_{Lipid\ 1}[s]\ - rt_{Lipid\ 2}[\mathrm{s}]$. *Values were first summarized per mouse (mean) and then summarized by calculation of mean and standard deviation of the twelve biological replicates. For 1-O-alk-1´-enyl/1-O-alkyl and +1 DB (∆1) negative ΔRT values resemble real RP-LC-MS conditions.*

| | **Lipid 1** | **Lipid 2** |
|---|---|---|
| **+2 CH2** | PC(34:0) | PC(32:0) |
| | PC(34:1) | PC(32:1) |
| | PC(34:2) | PC(32:2) |
| | PC(36:1) | PC(34:1) |
| | PC(36:2) | PC(34:2) |
| | PC(38:3) | PC(36:3) |
| | PE(34:0) | PE(32:0) |
| | PE(34:1) | PE(32:1) |
| | PE(34:2) | PE(32:2) |
| | PE(36:1) | PE(34:1) |
| | PE(36:2) | PE(34:2) |
| | PE(38:3) | PE(36:3) |
| **+1 DB (>Δ4)** | PC(32:1) | PC(32:0) |
| | PC(32:2) | PC(32:1) |
| | PC(34:1) | PC(34:0) |
| | PC(34:2) | PC(34:1) |
| | PC(34:3) | PC(34:2) |
| | PC(36:2) | PC(36:1) |
| | PC(36:3) | PC(36:2) |
| | PE(32:1) | PE(32:0) |
| | PE(32:2) | PE(32:1) |
| | PE(34:1) | PE(34:0) |
| | PE(34:2) | PE(34:1) |
| | PE(34:3) | PE(34:2) |
| | PE(36:2) | PE(36:1) |
| | PE(36:3) | PE(36:2) |
| | PE(P-34:1) | PE(P-34:0) |
| | PE(P-34:2) | PE(P-34:1) |
| | PE(P-34:3) | PE(P-34:2) |
| **+1 DB (Δ1)** | PC(P-36:1) | PC(O-36:1) |
| | PC(P-36:2) | PC(O-36:2) |
| | PC(P-36:4) | PC(O-36:4) |
| | PC(P-38:4) | PC(O-38:4) |
| | PC(P-38:6) | PC(O-38:6) |
| | PC(P-40:6) | PC(O-40:6) |
| | PC(P-40:8) | PC(O-40:8) |
| | PE(P-34:0) | PE(O-34:0) |
| | PE(P-34:1) | PE(O-34:1) |
| | PE(P-34:2) | PE(O-34:2) |
| | PE(P-36:1) | PE(O-36:1) |
| | PE(P-36:2) | PE(O-36:2) |
| | PE(P-36:5) | PE(O-36:5) |
| **1-O-alk-1´-enyl/1-O-alkyl** | PC(P-34:1) | PC(O-34:2) |
| | PC(P-36:1) | PC(O-36:2) |
| | PC(P-36:2) | PC(O-36:3) |
| | PC(P-36:3) | PC(O-36:4) |
| | PC(P-38:1) | PC(O-38:2) |
| | PC(P-38:5) | PC(O-38:6) |
| | PC(P-38:6) | PC(O-38:7) |
| | PC(P-40:7) | PC(O-40:8) |
| | PE(P-34:0) | PE(O-34:1) |
| | PE(P-34:1) | PE(O-34:2) |
| | PE(P-34:2) | PE(O-34:3) |
| | PE(P-35:1) | PE(O-35:2) |
| | PE(P-36:1) | PE(O-36:2) |
| | PE(P-36:4) | PE(O-36:5) |
| | PE(P-38:1) | PE(O-38:2) |

***Supplemental Table 2: HPLC method details**. Giving details about the different components and their respective settings regarding the used instrument Dionex Ultimate 3000 HPLC (Thermofisher Scientific Inc, Waltham, USA).*

| **HPLC Component** | **Applied settings** |
|---|---|
| Colum temperature | 50°C |
| Autosampler temperature | 10°C |
| Running solvent A | 10 mM ammonium formate and 0.2% formic acid in 6/4 (v/v) acetonitrile/H2O |
| Running solvent B | 10 mM ammonium formate and 0.2% formic acid in 9/1 (v/v) isopropanol/acetonitrile |
| Flow rate | 0.4 ml/min from 0 to 22 minutes<br>0.5 ml/min from 22 to 23 minutes<br>0.6 ml/min from 23 to 28 minutes<br>0.5 ml/min from 28 to 29 minutes<br>0.4 ml/min from 29 to 30 minutes |
| Gradient | 46% B for 2 min<br>46-72% B in 20 min<br>72-99% B in 1 min<br>99% B for 5 min<br>99-50% B in 1 min<br>46% B for 1 min |
| Sample amount | 10 µl |
| Washing volume | 200 µl |

***Supplemental Table 3: Mass spectrometric method details**. Giving details about the different components and their respective settings regarding the used instrument LTQ Velos MS (Thermo Fisher Scientific Inc., Waltham, USA).*

| **MS component** | **Setting** |
|---|---|
| Source | ESI negative mode |
| Capillary temperature | 275 °C |
| Source voltage | 3.8 kV |
| Scan rate | Normal |
| Activation type | CID |
| Full mass range | 460-1650 m/z |
| Isolation width | 1.5 |
| Normalized collision energy | 30/38 |
| Default charge state | 1 |
| Activation time | 10ms |
| MS2 mass range | 5-105% relative of parent mass |
| Number of micro scans Full MS | 3 |
| Number of micro scans MS2 | 5 |
| Dynamic exclusion | Enabled |
| Exclusion duration | 6 s |

***Supplemental Table 4: Peaklist for the integration of glycerophosphatidylcholines and glycerophosphatidylethanolamines**. Column 1: Lipid identifier (nomenclature described in [38]); Column 2: SubID for distinguishing between different peaks of the same isobaric species resulting from differing radyl combinations; Column 3: Respective retention times (in minutes); Column 4: Mass to charge ratios (m/z) of the lipids.*

| Lipid species | SubID | Retention time (min) | m/z |
|---|---|---|---|
| **PC(32:0)** | 1 | 4.51 | 778.729 |
| **PC(32:1)** | 1 | 4.23 | 776.645 |
| **PC(32:1)** | 2 | 3.88 | 776.620 |
| **PC(32:1)** | 3 | 4.05 | 776.620 |
| **PC(32:1)** | 4 | 3.69 | 776.586 |
| **PC(32:2)** | 1 | 2.99 | 774.561 |
| **PC(33:2)** | 1 | 3.40 | 788.565 |
| **PC(34:0)** | 1 | 5.78 | 806.561 |
| **PC(34:1)** | 1 | 4.72 | 804.658 |
| **PC(34:2)** | 1 | 3.86 | 802.511 |
| **PC(34:3)** | 1 | 3.16 | 800.610 |
| **PC(34:3)** | 2 | 3.40 | 800.565 |
| **PC(36:0)** | 1 | 7.31 | 834.658 |
| **PC(36:1)** | 1 | 6.03 | 832.606 |
| **PC(36:2)** | 1 | 5.01 | 830.584 |
| **PC(36:3)** | 1 | 4.11 | 828.547 |
| **PC(36:4)** | 1 | 3.73 | 826.479 |
| **PC(36:4)** | 2 | 3.30 | 826.511 |
| **PC(36:5)** | 1 | 3.08 | 824.494 |
| **PC(36:6)** | 1 | 2.62 | 822.496 |
| **PC(36:7)** | 1 | 2.61 | 820.513 |
| **PC(37:2)** | 1 | 5.69 | 844.607 |
| **PC(37:3)** | 1 | 4.72 | 842.591 |
| **PC(38:2)** | 1 | 6.51 | 858.779 |
| **PC(38:2)** | 2 | 6.16 | 858.722 |
| **PC(38:3)** | 1 | 5.61 | 856.607 |
| **PC(38:3)** | 2 | 5.39 | 856.607 |
| **PC(38:4)** | 1 | 4.91 | 854.618 |
| **PC(38:4)** | 2 | 4.55 | 854.583 |
| **PC(38:4)** | 3 | 4.35 | 854.548 |
| **PC(38:5)** | 1 | 4.18 | 852.591 |
| **PC(38:5)** | 2 | 3.97 | 852.574 |
| **PC(38:5)** | 3 | 3.76 | 852.591 |
| **PC(38:6)** | 1 | 3.47 | 850.478 |
| **PC(38:6)** | 2 | 3.18 | 850.495 |
| **PC(38:7)** | 1 | 2.84 | 848.536 |
| **PC(38:7)** | 2 | 2.63 | 848.532 |
| **PC(40:0)** | 1 | 10.83 | 890.685 |
| **PC(40:1)** | 1 | 9.12 | 888.647 |
| **PC(40:2)** | 1 | 7.84 | 886.654 |
| **PC(40:3)** | 1 | 6.92 | 884.638 |
| **PC(40:3)** | 2 | 6.62 | 884.651 |
| **PC(40:4)** | 1 | 6.32 | 882.610 |
| **PC(40:4)** | 2 | 5.82 | 882.641 |
| **PC(40:5)** | 1 | 5.41 | 880.691 |
| **PC(40:5)** | 2 | 5.03 | 880.630 |
| **PC(40:5)** | 3 | 4.81 | 880.630 |
| **PC(40:5)** | 4 | 4.14 | 880.630 |
| **PC(40:6)** | 1 | 4.53 | 878.712 |
| **PC(40:6)** | 2 | 4.13 | 878.527 |
| **PC(40:7)** | 1 | 3.62 | 876.610 |
| **PC(40:7)** | 2 | 3.36 | 876.610 |
| **PC(40:7)** | 3 | 3.18 | 876.610 |
| **PC(40:8)** | 1 | 2.91 | 874.569 |
| **PC(40:9)** | 1 | 2.44 | 872.526 |
| **PC(41:4)** | 1 | 5.60 | 868.615 |
| **PC(O-32:0)** | 1 | 5.30 | 764.664 |
| **PC(O-32:1)** | 1 | 4.35 | 762.629 |
| **PC(O-34:2)** | 1 | 4.64 | 788.599 |
| **PC(O-36:0)** | 1 | 6.25 | 820.576 |
| **PC(O-36:0)** | 2 | 5.06 | 820.614 |
| **PC(O-36:1)** | 1 | 7.23 | 818.532 |
| **PC(O-36:2)** | 1 | 5.94 | 816.612 |
| **PC(O-36:3)** | 1 | 4.84 | 814.623 |
| **PC(O-36:4)** | 1 | 4.50 | 812.583 |
| **PC(O-38:1)** | 1 | 7.59 | 846.662 |
| **PC(O-38:2)** | 1 | 7.52 | 844.677 |
| **PC(O-38:4)** | 1 | 5.75 | 840.634 |
| **PC(O-38:4)** | 2 | 4.36 | 840.634 |
| **PC(O-38:5)** | 1 | 4.63 | 838.586 |
| **PC(O-38:5)** | 2 | 4.40 | 838.681 |
| **PC(O-38:6)** | 1 | 4.08 | 836.609 |
| **PC(O-38:7)** | 1 | 3.25 | 834.584 |
| **PC(O-40:6)** | 1 | 5.41 | 864.632 |
| **PC(O-40:6)** | 2 | 5.26 | 864.637 |
| **PC(O-40:7)** | 1 | 4.26 | 862.611 |
| **PC(O-40:8)** | 1 | 3.39 | 860.591 |
| **PC(P-32:0)** | 1 | 5.18 | 762.592 |
| **PC(P-34:1)** | 1 | 5.49 | 788.522 |
| **PC(P-36:1)** | 1 | 6.97 | 816.612 |
| **PC(P-36:2)** | 1 | 5.73 | 814.596 |
| **PC(P-36:3)** | 1 | 5.28 | 812.581 |

| | | | |
|---|---|---|---|
| **PC(P-36:4)** | 1 | 4.36 | 810.490 |
| **PC(P-36:5)** | 1 | 3.09 | 808.448 |
| **PC(P-36:6)** | 1 | 2.50 | 806.504 |
| **PC(P-38:1)** | 1 | 7.81 | 844.671 |
| **PC(P-38:2)** | 1 | 5.53 | 842.645 |
| **PC(P-38:4)** | 1 | 5.54 | 838.662 |
| **PC(P-38:5)** | 1 | 4.75 | 836.581 |
| **PC(P-38:6)** | 1 | 4.02 | 834.608 |
| **PC(P-38:7)** | 1 | 3.34 | 832.549 |
| **PC(P-40:6)** | 1 | 5.11 | 862.566 |
| **PC(P-40:7)** | 1 | 4.12 | 860.618 |
| **PC(P-40:8)** | 1 | 3.29 | 858.664 |
| **PE(32:0)** | 1 | 4.83 | 690.540 |
| **PE(32:1)** | 1 | 3.98 | 688.510 |
| **PE(32:2)** | 1 | 3.19 | 686.460 |
| **PE(34:0)** | 1 | 6.17 | 718.517 |
| **PE(34:1)** | 1 | 5.11 | 716.764 |
| **PE(34:2)** | 1 | 4.17 | 714.621 |
| **PE(34:3)** | 1 | 3.38 | 712.576 |
| **PE(34:3)** | 2 | 3.65 | 712.478 |
| **PE(34:4)** | 1 | 3.09 | 710.627 |
| **PE(35:2)** | 1 | 4.76 | 728.608 |
| **PE(36:1)** | 1 | 6.53 | 744.540 |
| **PE(36:2)** | 1 | 5.45 | 742.507 |
| **PE(36:3)** | 1 | 4.42 | 740.478 |
| **PE(36:3)** | 2 | 4.81 | 740.559 |
| **PE(36:4)** | 1 | 4.07 | 738.482 |
| **PE(36:4)** | 2 | 3.58 | 738.450 |
| **PE(36:5)** | 1 | 3.25 | 736.479 |
| **PE(36:5)** | 2 | 2.89 | 736.519 |
| **PE(37:1)** | 1 | 7.23 | 758.576 |
| **PE(37:2)** | 1 | 6.13 | 756.629 |
| **PE(38:1)** | 1 | 8.17 | 772.621 |
| **PE(38:1)** | 2 | 7.97 | 772.657 |
| **PE(38:2)** | 1 | 6.95 | 770.564 |
| **PE(38:2)** | 2 | 6.67 | 770.617 |
| **PE(38:3)** | 1 | 6.28 | 768.533 |
| **PE(38:3)** | 2 | 5.97 | 768.533 |
| **PE(38:3)** | 3 | 5.78 | 768.560 |
| **PE(38:4)** | 1 | 5.27 | 766.503 |
| **PE(38:4)** | 2 | 4.88 | 766.476 |
| **PE(38:4)** | 3 | 4.73 | 766.503 |
| **PE(38:5)** | 1 | 4.27 | 764.532 |
| **PE(38:5)** | 2 | 3.97 | 764.583 |
| **PE(38:5)** | 3 | 4.56 | 764.532 |
| **PE(38:6)** | 1 | 3.70 | 762.509 |
| **PE(38:6)** | 2 | 3.42 | 762.476 |
| **PE(38:7)** | 1 | 3.97 | 760.458 |

| | | | |
|---|---|---|---|
| **PE(38:7)** | 2 | 3.65 | 760.501 |
| **PE(38:7)** | 3 | 3.49 | 760.487 |
| **PE(39:4)** | 1 | 5.97 | 780.604 |
| **PE(40:1)** | 1 | 9.99 | 800.586 |
| **PE(40:1)** | 2 | 10.46 | 800.567 |
| **PE(40:2)** | 1 | 8.36 | 798.594 |
| **PE(40:3)** | 1 | 7.35 | 796.720 |
| **PE(40:4)** | 1 | 6.76 | 794.631 |
| **PE(40:4)** | 2 | 6.21 | 794.559 |
| **PE(40:5)** | 1 | 5.82 | 792.594 |
| **PE(40:5)** | 2 | 5.42 | 792.671 |
| **PE(40:5)** | 3 | 5.10 | 792.550 |
| **PE(40:6)** | 1 | 4.86 | 790.583 |
| **PE(40:6)** | 2 | 4.21 | 790.625 |
| **PE(40:6)** | 3 | 3.51 | 790.550 |
| **PE(40:7)** | 1 | 3.90 | 788.509 |
| **PE(40:8)** | 1 | 3.11 | 786.512 |
| **PE(40:9)** | 1 | 2.60 | 784.543 |
| **PE(42:2)** | 1 | 10.01 | 826.608 |
| **PE(42:2)** | 2 | 9.67 | 826.608 |
| **PE(42:3)** | 1 | 8.69 | 824.720 |
| **PE(42:4)** | 1 | 7.50 | 822.705 |
| **PE(42:5)** | 1 | 6.34 | 820.631 |
| **PE(42:6)** | 1 | 5.37 | 818.660 |
| **PE(42:7)** | 1 | 4.46 | 816.680 |
| **PE(O-34:0)** | 1 | 7.30 | 704.577 |
| **PE(O-34:1)** | 1 | 6.03 | 702.662 |
| **PE(O-34:2)** | 1 | 4.99 | 700.679 |
| **PE(O-34:3)** | 1 | 4.04 | 698.658 |
| **PE(O-34:3)** | 2 | 4.37 | 698.568 |
| **PE(O-34:4)** | 1 | 3.69 | 696.591 |
| **PE(O-35:1)** | 1 | 6.75 | 716.764 |
| **PE(O-35:2)** | 1 | 5.63 | 714.621 |
| **PE(O-35:4)** | 1 | 4.24 | 710.579 |
| **PE(O-36:1)** | 1 | 7.71 | 730.631 |
| **PE(O-36:1)** | 2 | 7.46 | 730.562 |
| **PE(O-36:2)** | 1 | 6.37 | 728.621 |
| **PE(O-36:3)** | 1 | 5.83 | 726.594 |
| **PE(O-36:3)** | 2 | 5.62 | 726.533 |
| **PE(O-36:3)** | 3 | 5.36 | 726.594 |
| **PE(O-36:3)** | 4 | 5.12 | 726.628 |
| **PE(O-36:4)** | 1 | 4.83 | 724.550 |
| **PE(O-36:5)** | 1 | 4.11 | 722.444 |
| **PE(O-36:5)** | 2 | 3.88 | 722.479 |
| **PE(O-36:6)** | 1 | 3.33 | 720.446 |
| **PE(O-37:4)** | 1 | 5.51 | 738.570 |
| **PE(O-37:4)** | 2 | 5.37 | 738.570 |
| **PE(O-37:6)** | 1 | 3.78 | 734.522 |

| | | | |
|---|---|---|---|
| **PE(O-38:0)** | 1 | 10.14 | 760.658 |
| **PE(O-38:0)** | 2 | 10.63 | 760.597 |
| **PE(O-38:1)** | 1 | 9.20 | 758.587 |
| **PE(O-38:2)** | 1 | 8.13 | 756.600 |
| **PE(O-38:2)** | 2 | 7.80 | 756.608 |
| **PE(O-38:3)** | 1 | 7.44 | 754.593 |
| **PE(O-38:3)** | 2 | 6.83 | 754.610 |
| **PE(O-38:4)** | 1 | 6.20 | 752.569 |
| **PE(O-38:4)** | 2 | 5.73 | 752.587 |
| **PE(O-38:5)** | 1 | 5.37 | 750.615 |
| **PE(O-38:5)** | 2 | 5.10 | 750.625 |
| **PE(O-38:5)** | 3 | 4.80 | 750.605 |
| **PE(O-38:6)** | 1 | 4.42 | 748.574 |
| **PE(O-38:7)** | 1 | 3.56 | 746.549 |
| **PE(O-39:5)** | 1 | 7.06 | 766.619 |
| **PE(O-39:7)** | 1 | 5.01 | 762.604 |
| **PE(O-40:1)** | 1 | 11.13 | 786.805 |
| **PE(O-40:2)** | 1 | 9.96 | 784.664 |
| **PE(O-40:3)** | 1 | 8.56 | 782.576 |
| **PE(O-40:3)** | 2 | 8.34 | 782.584 |
| **PE(O-40:4)** | 1 | 7.95 | 780.588 |
| **PE(O-40:4)** | 2 | 7.31 | 780.633 |
| **PE(O-40:5)** | 1 | 6.81 | 778.663 |
| **PE(O-40:5)** | 2 | 6.36 | 778.618 |
| **PE(O-40:5)** | 3 | 6.13 | 778.618 |
| **PE(O-40:6)** | 1 | 5.69 | 776.668 |
| **PE(O-40:7)** | 1 | 4.59 | 774.612 |
| **PE(O-40:8)** | 1 | 3.69 | 772.544 |
| **PE(O-40:9)** | 1 | 2.99 | 770.544 |
| **PE(O-42:10)** | 1 | 4.80 | 796.600 |
| **PE(O-42:12)** | 1 | 4.44 | 792.600 |
| **PE(O-42:6)** | 1 | 7.30 | 804.703 |
| **PE(O-42:7)** | 1 | 5.86 | 802.612 |
| **PE(P-34:0)** | 1 | 7.14 | 702.454 |
| **PE(P-34:1)** | 1 | 5.88 | 700.599 |
| **PE(P-34:2)** | 1 | 4.84 | 698.570 |
| **PE(P-34:3)** | 1 | 4.21 | 696.398 |
| **PE(P-34:3)** | 2 | 3.99 | 696.528 |
| **PE(P-34:4)** | 1 | 3.57 | 694.335 |
| **PE(P-35:1)** | 1 | 6.59 | 714.329 |
| **PE(P-35:2)** | 1 | 5.49 | 712.673 |
| **PE(P-35:4)** | 1 | 4.12 | 708.436 |
| **PE(P-36:1)** | 1 | 7.38 | 728.620 |
| **PE(P-36:2)** | 1 | 6.20 | 726.500 |
| **PE(P-36:3)** | 1 | 5.63 | 724.515 |
| **PE(P-36:3)** | 2 | 5.41 | 724.515 |
| **PE(P-36:3)** | 3 | 5.13 | 724.635 |
| **PE(P-36:4)** | 1 | 4.67 | 722.479 |
| **PE(P-36:5)** | 1 | 3.82 | 720.529 |
| **PE(P-36:6)** | 1 | 3.25 | 718.536 |
| **PE(P-37:4)** | 1 | 5.35 | 736.525 |
| **PE(P-38:1)** | 1 | 9.02 | 756.602 |
| **PE(P-38:2)** | 1 | 7.93 | 754.512 |
| **PE(P-38:2)** | 2 | 7.61 | 754.646 |
| **PE(P-38:3)** | 1 | 7.16 | 752.578 |
| **PE(P-38:3)** | 2 | 6.62 | 752.467 |
| **PE(P-38:4)** | 1 | 6.01 | 750.615 |
| **PE(P-38:4)** | 2 | 5.59 | 750.605 |
| **PE(P-38:5)** | 1 | 5.17 | 748.502 |
| **PE(P-38:5)** | 2 | 4.95 | 748.553 |
| **PE(P-38:5)** | 3 | 4.76 | 748.584 |
| **PE(P-38:6)** | 1 | 4.25 | 746.474 |
| **PE(P-38:6)** | 2 | 3.97 | 746.535 |
| **PE(P-38:7)** | 1 | 3.52 | 744.498 |
| **PE(P-39:5)** | 1 | 6.80 | 764.569 |
| **PE(P-39:7)** | 1 | 4.83 | 760.487 |
| **PE(P-40:1)** | 1 | 10.91 | 784.727 |
| **PE(P-40:2)** | 1 | 9.19 | 782.690 |
| **PE(P-40:3)** | 1 | 8.33 | 780.556 |
| **PE(P-40:4)** | 1 | 7.73 | 778.648 |
| **PE(P-40:4)** | 2 | 7.17 | 778.633 |
| **PE(P-40:5)** | 1 | 6.55 | 776.628 |
| **PE(P-40:5)** | 2 | 6.20 | 776.655 |
| **PE(P-40:5)** | 3 | 5.98 | 776.664 |
| **PE(P-40:6)** | 1 | 5.53 | 774.629 |
| **PE(P-40:7)** | 1 | 4.49 | 772.499 |
| **PE(P-40:8)** | 1 | 3.60 | 770.620 |
| **PE(P-40:9)** | 1 | 2.93 | 768.510 |
| **PE(P-42:10)** | 1 | 4.24 | 794.528 |
| **PE(P-42:6)** | 1 | 7.09 | 802.578 |
| **PE(P-42:7)** | 1 | 5.67 | 800.618 |
| **PE(P-42:8)** | 1 | 4.60 | 798.526 |

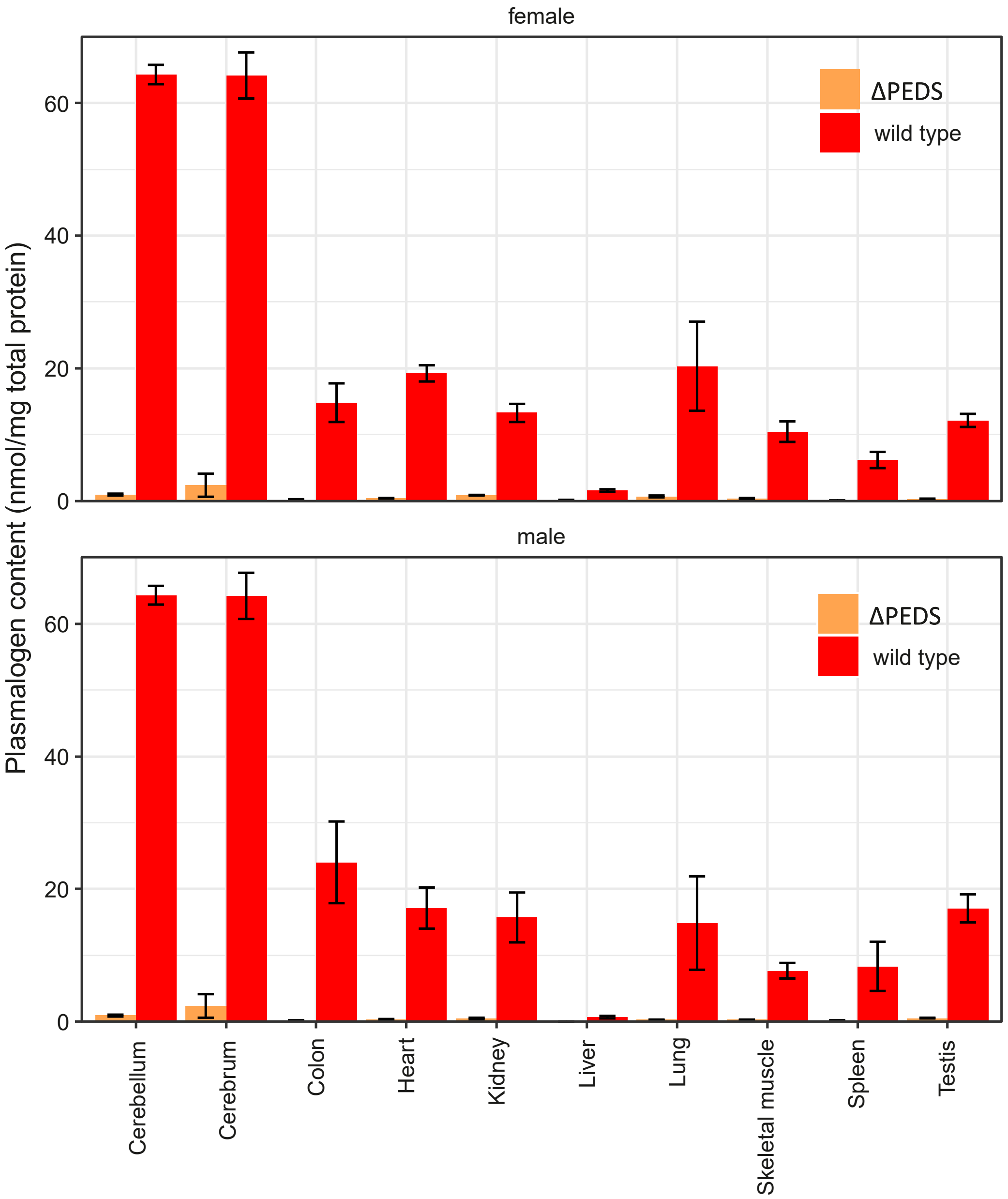


***Supplemental Figure 1: Plasmalogen content per mg total protein.*** *Measured using the plasmalogen analysis method reported in [29]. Data visualized from values in [5].*
